# Conjugation structures plasmid populations through host-lineage restriction

**DOI:** 10.64898/2026.02.19.706745

**Authors:** William Matlock, R. Craig MacLean

## Abstract

Conjugation mediates plasmid transfer between bacterial species, driving the horizontal spread of traits like antibiotic resistance. However, genomic and experimental evidence indicates that many conjugative plasmids are restricted to particular host lineages despite broad theoretical host ranges. Using 4,281 plasmids from an epidemiologically coherent sample of 1,739 *Escherichia coli* bloodstream infection isolates, we quantify the host-lineage associations of 30 plasmid backbones and assess how these associations structure plasmid co-occurrence. To achieve this, we develop a novel Bayesian modelling framework that separates genuine backbone–backbone associations from patterns arising due to shared host ancestry or abundance. First, we find that conjugative backbones exhibit stronger host-lineage restriction than mobilisable backbones, and that restriction increases with plasmid size independently of mobility class. Consistent with this pattern, host-lineage restricted backbones demonstrate fewer independent acquisitions across the host phylogeny in ancestral state reconstructions. Comparison with global plasmid diversity shows that the most host-lineage restricted backbones remain restricted beyond the studied population, whereas the least restricted backbones span a mean of seven host species. Next, after accounting for host phylogeny and abundance, we find that two thirds of backbone pairs show no strong association or avoidance; however, backbones sharing host lineages co-occur more frequently than expected. Lastly, we identify a clique of strongly associated mobilisable backbones that appear to exploit a shared set of lineage-restricted conjugative partners. A mathematical model demonstrates that host-lineage restriction of conjugative backbones, together with specificity in conjugative–mobilisable transfer, is sufficient to generate clustering among mobilisable plasmids. Collectively, our findings reframe conjugation as a mechanism that promotes within-lineage persistence and shapes plasmid community structure, with potentially important consequences for the accumulation of resistance and virulence determinants.

## Introduction

Plasmids are mobile genetic elements ubiquitous across bacteria, and their transfer between hosts is a major driver of horizontal gene transfer and adaptive evolution, particularly in the context of antibiotic resistance [1]. Plasmids move between hosts either by encoding a complete conjugative apparatus or, in the case of mobilisable plasmids, by encoding only partial transfer functions and piggybacking on co-occurring conjugative elements [2]. As plasmids disseminate, cargo genes may be gained or lost while a conserved backbone encoding essential replication and maintenance functions is retained, allowing plasmid lineages to be defined despite substantial gene-content variation [3]. Understanding plasmid evolution therefore requires distinguishing between the capacity for transfer and the realised patterns of persistence and spread observed at the population level.

Genomic surveillance [4, 5, 6] and large sequence database analyses [7, 8] have shown that some backbones are widespread, whereas others preferentially associate with particular bacterial host lineages and can persist there for centuries [9]. These patterns suggest that, despite their theoretical host range, many plasmids are effectively limited to a subset of bacteria, indicating that recipient lineage plays a central role in shaping plasmid distribution.

Experimental studies provide evidence that plasmid transfer and persistence are often host-lineage specific. Plasmids preferentially conjugate to closely related hosts [10], in part through donor-encoded mating pair stabilisation proteins that restrict plasmid transfer to recipients carrying compatible outer-membrane receptors [11, 12, 13]. Following this initial transfer, long-term plasmid persistence is shaped by strain-specific epistasis [14]. For example, the carbapenemase plasmid pOXA-48 encodes a transcriptional regulator that alters chromosomal expression in *Klebsiella* and *Citrobacter*, producing a fitness benefit that reinforces lineage-specific persistence [15]. In addition, co-occurring plasmids can exhibit positive epistasis for fitness, reducing the combined fitness cost of carriage and thereby promoting long-term persistence under certain conditions [16]. However, such effects are not universal and vary between host lineages, with broader genetic and ecological context shaping outcomes [17]. Together, these findings suggest that apparent plasmid associations in population-scale data may reflect shared host compatibility and evolved mobility strategies rather than direct functional interactions alone.

Plasmid host-lineage restriction implies that analyses based on raw plasmid co-occurrence counts can be misleading. Consider the following scenarios involving two backbones. **(1)** They appear together frequently simply because they are both common in the sample; their overlap reflects abundance, not a functional association. **(2)** They never appear together even though they are compatible, creating the false impression that they avoid one another, when in reality, they just inhabit different host lineages. **(3)** They seem strongly linked simply because they live in the same host lineages; their co-occurrence is driven by shared hosts rather than any functional relationship. **(4)** They are found together more often than expected once abundance and shared hosts are taken into account, reflecting genuine association.

To address these issues, we developed a Bayesian hierarchical phylogenetic framework to model the strength of plasmid backbone–host lineage associations and the extent to which this influenced patterns in backbone co-occurrence. Shared ancestry creates non-independence among isolates, so observed patterns in backbone presence–absence may reflect lineage history. By explicitly modelling phylogenetic covariance, we partitioned variation attributable to ancestry from that arising from genuine backbone–backbone associations and avoidances [18]. We then used this model to reanalyse the NORM collection, an epidemiologically coherent population of *n* = 1,739 human bloodstream infection *Escherichia coli* isolates sampled between 2002-2017 in Norway [9]. In total, we analysed *n* = 4,281 plasmids resolved into *n* = 30 putative backbones.

## Materials and Methods

### Dataset curation

We first retrieved the metadata for the *n* = 4, 569 plasmids, including plasmid type (pT), predicted mobility (determined via MOB-suite [19]), and host sequence type (ST). We also retrieved the host chromosomal phylogeny, which was inferred in IQ-TREE using a core SNP alignment made by mapping short-reads to the EC958 reference genome [20].

pTs were defined in the original study by clustering complete plasmid sequences based on backbone similarity using the tool mge-cluster, followed by network-based community detection (Louvain method) to group plasmids that consistently co-clustered across multiple runs [21]. Each pT therefore represents a distinct plasmid backbone lineage. Plasmids were assigned one of *n* = 30 pTs. We found *n* = 236 instances of the same pT duplicated at least once in a genome, potentially caused by misassembly or genuine heterogeneity. In these cases, a random representative was chosen.

The NORM long-read cohort used a two-stage sampling strategy, first capturing overall diversity, then exhaustively including isolates from four major ExPEC lineages (ST69, ST73, ST95, and ST131). Although this oversampling gives these lineages greater influence on the global patterns, the hierarchical model’s partial pooling (described subsequently) stabilises estimates for underrepresented STs, preventing them from being dominated by the more densely sampled groups.

### Model formulation

We modelled the presence or absence of each pT across host tips using a Bayesian hierarchical logistic regression that jointly captured phylogenetic dependence among hosts and correlation among pTs. For tips *i* ∈ {1, …, *N*} and pTs *b* ∈ {1, …, *B*}, we modelled the presence of a pT as:

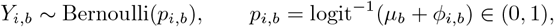

where *µ*_*b*_ is the pT-specific intercept and *ϕ*_*i,b*_ is the phylogenetic effect of host tip *i* on pT *b*. This formulation separated overall pT prevalence (*µ*_*b*_) from phylogenetically structured variation (*ϕ*_*i,b*_). The hierarchical framework mitigated uneven sampling by shrinking noisy estimates toward the global mean, enabling rare pT–host and pT–pT combinations to be interpreted alongside well-sampled groups with greater reliability.

### Intercept hierarchy

Intercepts were drawn from a multivariate normal distribution with shared mean, hierarchical scale, and correlated deviations:

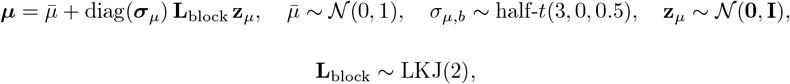

where **L**_block_ is the Cholesky factor of the pT correlation matrix. This allowed intercepts for related pTs to covary while remaining non-centred for efficient sampling. We used LKJ(2) for the correlation matrices to slightly shrink correlations toward zero. This reduced weak or noisy correlations, helping the model highlight stronger, more robust associations between pTs.

### Phylogenetic effects

Host phylogeny was represented by a symmetrised and standardised covariance matrix whose mean eigenvalue was scaled to one. We retained the top *J* = 245 eigenvectors and eigenvalues, explaining at least 99% of total variance, balancing computational efficiency and model fidelity. The phylogenetic effects were modelled in this reduced eigenbasis as:

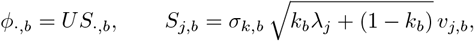

where *U* and *λ* are the retained eigenvectors and eigenvalues, *σ*_*k,b*_ ≥ 0 is the pT-specific phylogenetic standard deviation, and *k*_*b*_ ∈ (0, 1) controls the degree of phylogenetic signal, interpolating between a fully phylogenetic (*k*_*b*_ = 1) and independent (*k*_*b*_ = 0) process. The latent factors *v*_*j,b*_ induced a consistent pT correlation structure in both the intercepts and the phylogenetic terms:

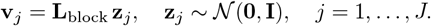

Each pT column of *ϕ*_·,*b*_ was mean-centred to prevent confounding with *µ*_*b*_:

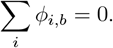

Weakly informative priors were used for the phylogenetic scales and signals:

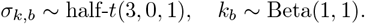

Under this standardisation, *σ*_*k,b*_ approximated the marginal standard deviation of the logit-scale phylogenetic effect *ϕ*_·,*b*_. Thus, logit^−1^(±*σ*_*k,b*_) provided a heuristic range of tip-level presence probabilities corresponding to a ±1 SD phylogenetic deviation (for *µ*_*b*_ = 0).

### Modelling pT co-occurrence

Each pT pair (*i, j*) was classified along two axes: phylogenetic (Φ_*ij*_) and residual (*R*_*ij*_) co-occurrence. We computed these metrics for each posterior draw *d* to obtain full posterior distributions. The phylogenetic axis Φ_*ij*_ was computed as the correlation of the latent effects across tips within each draw:

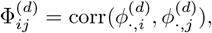

where 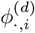 is the vector of latent effects for pT *i* in draw *d*. The residual axis *R*_*ij*_ was computed from the latent deviations after accounting for the average latent effect per host tip. For each draw, we centered the latent matrix by removing row means and computed the correlation of the resulting residuals:

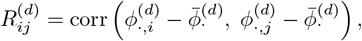

where 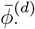 represents the vector of row means for draw *d*. The global relationship between phylogenetic and residual co-occurrence was then quantified by calculating the Pearson correlation between these point estimates across all unique pT pairs. Posterior certainty was quantified by the signed tail probability (STP):

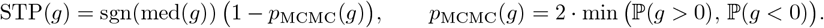

pT pairs with STP ≥ 0.95 were classified as positive, STP ≤ −0.95 as negative, and neutral otherwise. In addition, we required at least one observation for a pair to be classified as positive. The model still produced estimates for unobserved pairs by borrowing information across all pTs and phylogenetic structure; these can be interpreted as extrapolations.

### Model implementation and convergence

The model was implemented in Stan (see `model.stan`) and fitted from R via `cmdstanr` using four chains, each with 2,000 warmup and 2,000 sampling iterations [22, 23]. Model diagnostics indicated strong performance for top-level parameters (*k*_*b*_, *σ*_*k,b*_, *µ*_*b*_, 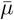, *σ*_*µ*_, and Φ_*ij*_) with 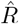 median = 1.00, range = 1.00–1.01; bulk effective sample size (ESS) median = 4635.2, range = 746.9–12,921.9; tail ESS median = 5,903, range = 1,579–7,755. Trace plots for all parameters were also visually inspected for convergence.

### Posterior predictive checks

To assess model fit, we performed posterior predictive checks by simulating datasets *y*^rep^ from each posterior draw of the predicted probabilities *p*. The observed mean plasmid presence (0.082) closely matched the posterior predictive mean (0.082), with a Bayesian *p*-value of 0.53, indicating that the observed mean was typical under the model. The observed standard deviation was also well captured (posterior predictive sd = 0.275; observed sd = 0.274; Bayesian *p*-value = 0.52). At the pT level, all observed frequencies fell within the 95% posterior predictive intervals, showing that the model accurately reproduced both overall and pT-specific patterns.

### Contextualising pTs in PLSDB diversity

We first downloaded *n* = 72, 556 plasmids and linked metadata from PLSDB (v. 2024_05_31_v2). Next, we used sourmash (v. 4.9.4; default parameters except k=21) to calculate the similarity of the NORM plasmids to the PLSDB plasmids. To call a hit, we required a 0.9 Jaccard similarity of *k*-mers. PLSDB metadata provided the host taxonomic classifications via NCBI. To avoid self-comparisons, we filtered out any PLSDB plasmids with matching nucleotide accessions to those in the NORM BioProjects (PRJEB45354 and PRJEB57633). We found no hits for pT 20-1. A total of 33 PLSDB plasmids mapped to multiple pTs. Most of these (30/33) were combinations of 2-1, 2-2, and 2-3, which were related F-type plasmids. Nonetheless, all 33 instances originated from *E. coli* isolates, so did not impact downstream analyses.

Plasmids ranged approximately from 1 kbp to 200 kbp, meaning that for `k=21` they contained 980 to 199,980 *k*-mers, respectively. With a Jaccard threshold of 0.9, roughly 10% of *k*-mers could differ between pairs and still be considered a match, corresponding to ∼100 and ∼20,000 *k*-mer differences for a 1 kbp and 200 kbp plasmid, respectively. This is approximately 10% because sourmash uses a MinFrac sketch, rather than working with *k*-mers directly. Since smaller plasmids are generally more conserved than larger plasmids, this threshold was suitable. Because PLSDB reflects sequenced diversity and is biased towards common pathogenic species such as *E. coli*, not detecting a pT in a given species did not imply that it never resides there (absence of evidence rather than evidence of absence).

The Poisson regression was implemented using the `brms` library in R [24]. Convergence was strong, with 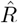 = 1.00, bulk ESS > 8, 000, and tail ESS > 10, 000. To calculate the expected number of species, we back-transformed the linear predictor from the log scale using the inverse link function:

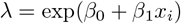

where *β*_0_ is the intercept, *β*_1_ is the coefficient for host-lineage restriction probability *P* (*k* ≥ 0.7).

Increasing the Jaccard similarity threshold to a more conservative 0.95, the model parameter estimates changed minimally and remained consistent with the previous 95% CIs (new intercept = 1.72 [95% CI = 1.40–2.04] and new slope = -1.51 [95% CI = -2.28 to -0.77]).

### Ancestral state reconstruction

We ran SimPhyNI (v. 1.0.0) [25] with default parameters except `–min_prev 0 –max_prev 1`, which utilises PastML (v. 1.9.50) [26] for ancestral state reconstruction. PastML was configured with default parameters except`––prediction_method JOINT -m F81`. This model reconstructs the most likely global combination of states across all internal nodes simultaneously, assuming uniform transition rates while accounting for unequal equilibrium state frequencies. Gain-bias 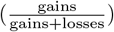was calculated directly from the PastML output. The binomial regressions were implemented using the `brms` library in R [24]. Convergence was strong across both models, with all 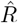 = 1.00, bulk ESS > 2,000, and tail ESS > 2,000.

SimPhyNI’s primary use is to detect instances of convergent association or avoidance. We compared these findings to our model outputs in Supplementary File 1.

### Modelling mobilisable plasmid clustering

We let there be *n*_hosts_ = 1,000 hosts, *n*_lineages_ = 5 host lineages, *n*_conj_ = 5 conjugative backbones, and *n*_mob_ = 5 mobilisable backbones. We then defined a mobilisation matrix 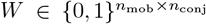, where *W*_*ij*_ = 1 if mobilisable plasmid *i* could use conjugative plasmid *j* for transfer. The matrix was constructed to reflect our observed patterns: some conjugative plasmids mobilised multiple mobilisable plasmids, and some mobilisable plasmids shared overlapping conjugative partners. For each conjugative plasmid *j*, we assigned a set of host lineages *L*_*j*_ ⊆ {1, …, *n*_lineages_} it could occupy. For host *k*, we let *h*_*k*_ ∈ {1, …, *n*_lineages_} denote its lineage, and drew conjugative plasmid presence as

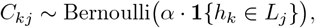

where *α* ∈ [0, 1] controlled host-lineage restriction. Mobilisable plasmids were then deterministically present if any linked conjugative plasmid was present:

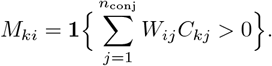

The co-occurrence matrix of mobilisable plasmids was

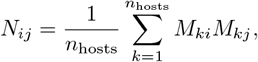

and within-cluster and between-cluster pairs were denoted by *P*_within_ and *P*_between_. The clustering score was computed as

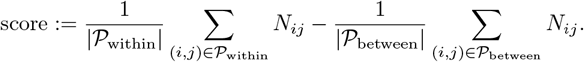

This model could equally be expressed as a directed acyclic graph *H* → *C*_*j*_ → *M*_*i*_, with host lineage *H* drawn first independently,

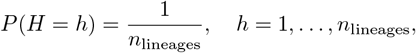

followed by independent conjugative plasmid draws conditional on *H*,

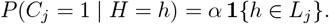

Mobilisable plasmids inherit this bias through compatibility:

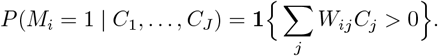

When *α* → 0, lineage bias vanished and *P* (*M*_*i*_ = 1, *M*_*k*_ = 1) ≈ *P* (*M*_*i*_ = 1)*P* (*M*_*k*_ = 1), leaving co-occurrence only from overlapping conjugative partners. Thus, co-occurrence arose entirely from shared conjugative partners and host-lineage restriction, without requiring additional dynamics.

## Results

### Plasmid backbones differed in strength of host-lineage associations

A plasmid backbone might prefer a specific host lineage, distribute widely, or exhibit an intermediate or more complex pattern. We started by resolving the *n* = 4,281 plasmids into *n* = 30 putative backbone groups using a sequence clustering approach, hereafter referred to as “plasmid types” (pTs; see Methods). These were previously characterised in [9]. Our modelling framework estimated pT-level parameters capturing both the strength of host-lineage restriction for each pT and the extent to which this restriction drove co-occurrence with other pTs. Figure 1a illustrates the conceptual workflow of the model. Comparison of the model to existing methodology is given in Supplementary File 1.

**Figure 1:**
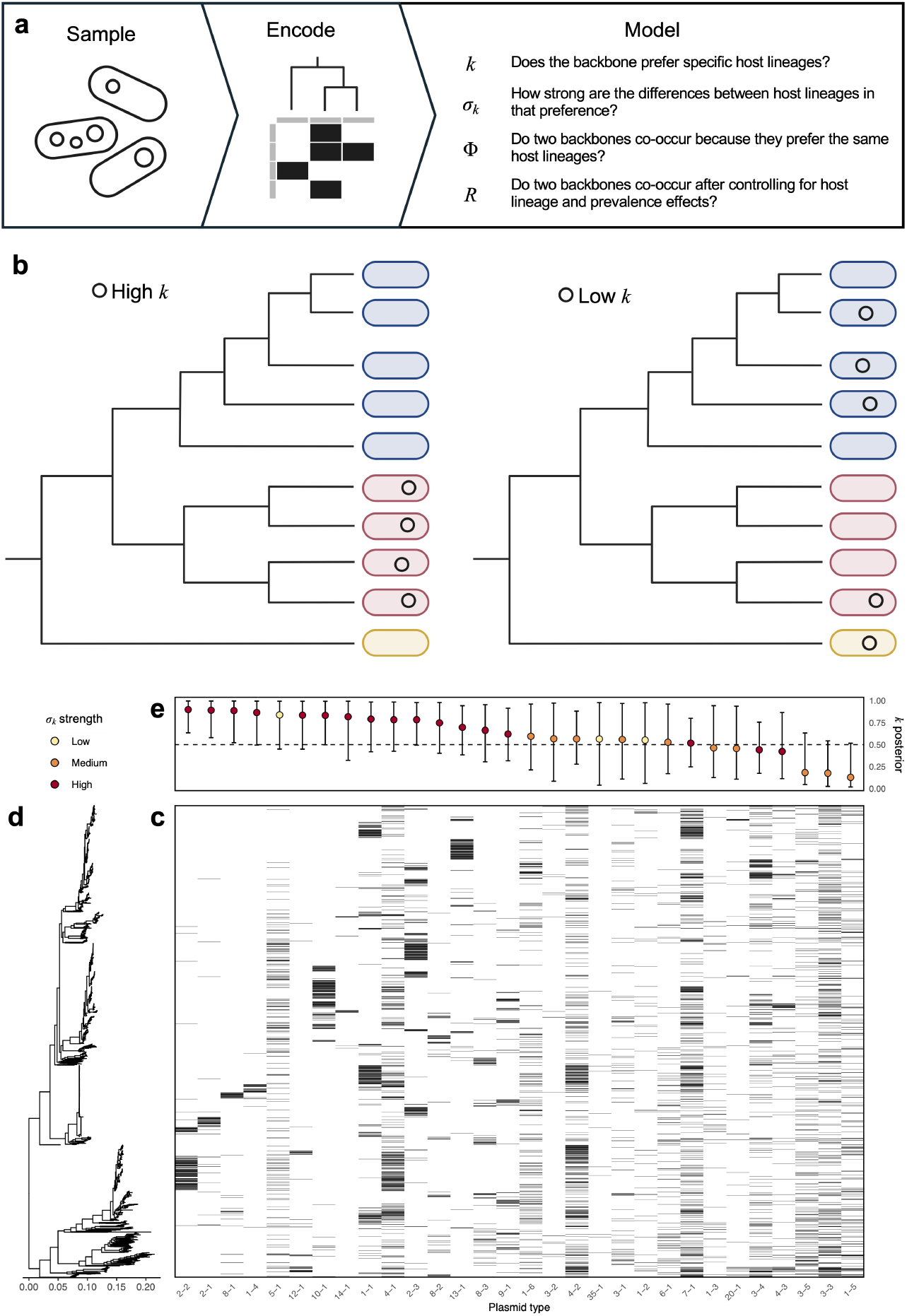
Modelling plasmid backbone host-lineage restriction. **(a)** Modelling workflow schematic. Sampled isolates are sequenced and assembled, then encoded as a host chromosomal phylogeny and a binary presence/absence matrix of plasmid backbones. The model then estimates parameters of interest for individual backbones and pairs of backbones (*k, σ*_*k*_, Φ, and *R*). **(b)** An example of a plasmid with a high or low *k* parameter. **(c)** Presence/absence matrix for all *n* = 30 plasmid types (pTs) with rows ordered by **(d)** host phylogeny and columns by **(e)** decreasing phylogenetic signal *k* posterior median (dots; lines indicate 95% credible intervals). Point colour encodes the median phylogenetic standard deviation (*σ*_*k*_), categorised as low (0–1.5), medium (1.5–3), and high (> 3). This parameter measures the absolute magnitude of the phylogenetic effect on the logit scale, where logit^−1^(± *σ*_*k*_) gives the probability that a tip’s presence or absence is determined by phylogeny alone: low shifts probabilities by ∼18–82%, medium by∼ 5–95%, and high is nearly deterministic (< 5% or > 95%).

Then, we quantified host-lineage associations by developing a paired measure: phylogenetic signal (*k*; 0–1), which asked how strongly variation in pT presence was structured by host relatedness (independent of pT frequency), and signal strength (*σ*_*k*_), which asked how large that phylogenetically structured variation was (see Figure 1b). At the extremes, taking high or low values, we obtain four possible classifications of host-lineage associations (Table 1).

**Table 1:**
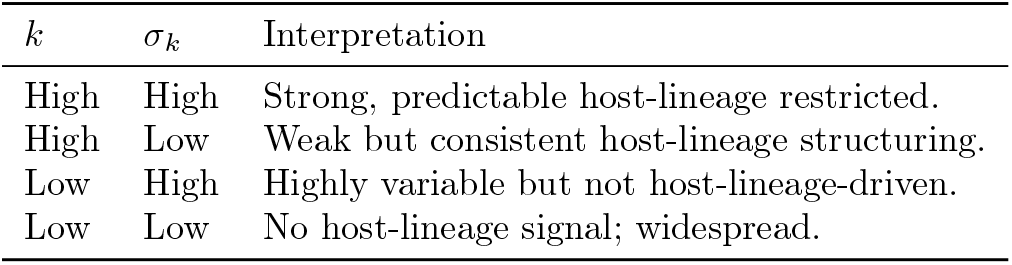
Interpretation of model parameters *k* and *σ*_*k*_.

Figure 1c shows the presence/absence matrix, ordered by host chromosomal phylogeny (rows; Figure 1d) and decreasing posterior median *k* (columns; Figure 1e). Note that since our approach was Bayesian, we estimated posterior distributions for *k* and *σ*_*k*_, not a single value. Therefore, to assess support for strong and weak host-lineage restriction, we calculated the posterior probabilities that *k* ≥ 0.7 (strong) and *k* ≤ 0.3 (weak), respectively. pTs showed variable evidence for phylogenetic structure, with a median P(*k* ≥ 0.7) of 0.34 (range = 0.00–0.94) and a median P(*k* ≤ 0.3) of 0.03 (range = 0.00–0.87), suggesting a spectrum from host-lineage restricted to widespread.

### Host-lineage restricted backbones remained restricted in global sample

While a plasmid backbone may be host-lineage restricted within our *E. coli* population, closely related plasmids sampled elsewhere could occur in other species. Conversely, a backbone that appeared unrestricted might still be confined to *E. coli* in other contexts, or extend across the entire Enterobacterales order.

First, we screened our plasmids for identical or near-identical matches in the *n* = 72,556 plasmid sequences in PLSDB, a lightly curated subset derived from NCBI, providing a snapshot of global sequenced plasmid diversity. Then, for each pT, we recorded the number of unique taxonomic representatives among the PLSDB hits. We found that some pTs had close relatives across multiple taxonomic ranks, from species to phyla (Figure 2a).

**Figure 2:**
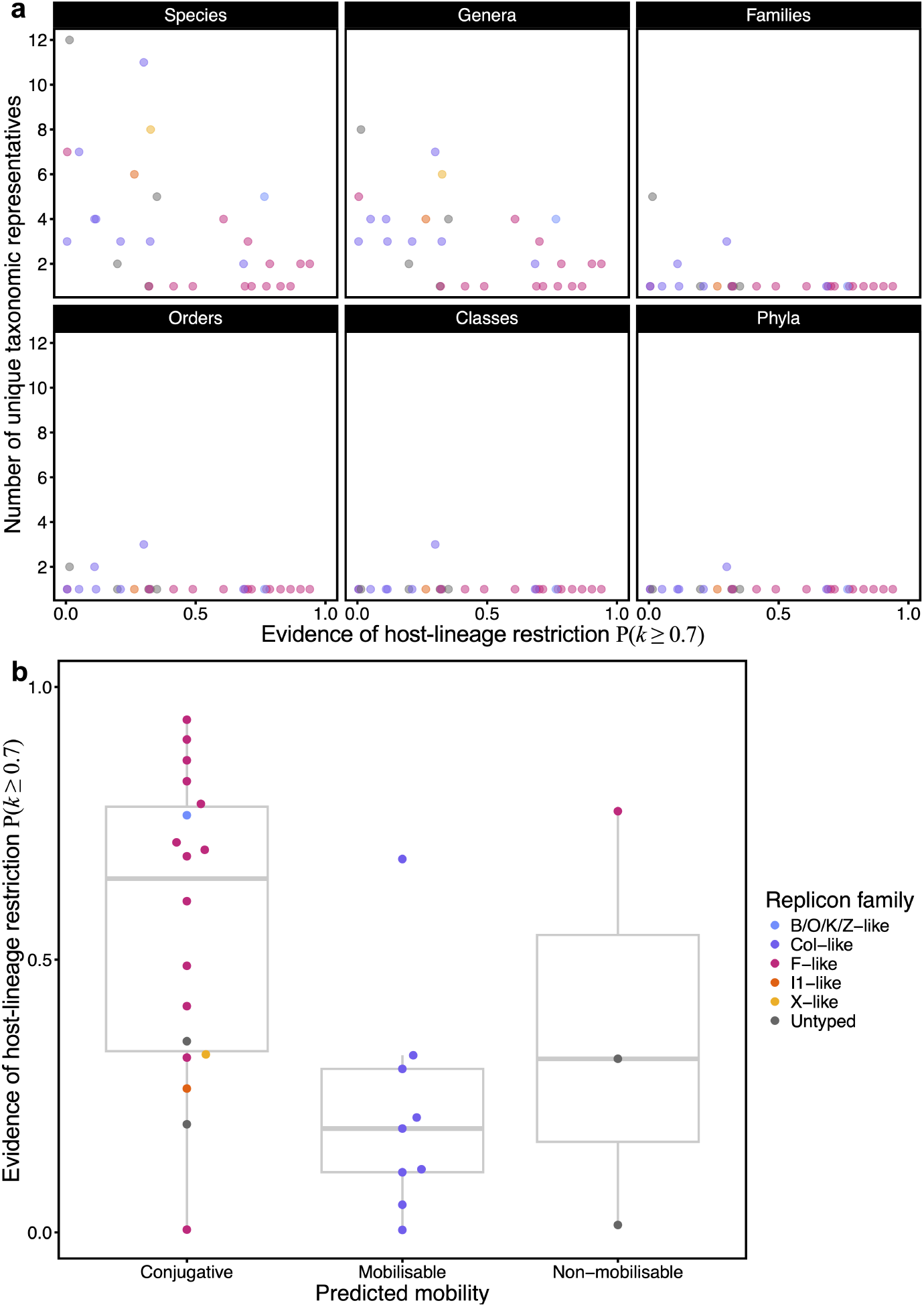
Host-lineage restriction across global diversity and predicted plasmid mobility. **(a)** Taxonomic rank distributions of plasmid type (pT) relatives in gobal sequenced diversity. Each panel shows a different taxonomic rank. The *x*-axes shows the probability that the pT was host lineage-restricted in the sample, determined via the model. The *y*-axes show the number of unique taxonomic rank representatives found in PLSDB by screening the NORM plasmids for identical or near-identical relatives, then mapping back to the respective pT. Note that pT 20-1 had no hits, so only 29 points are shown in each panel. **(b)** The *x*-axis shows the predicted mobility type for each pT. The *y*-axis shows the probability that the pT was host lineage-restricted. Colours across panels indicate the pT replicon family.

Next, we predicted pT species host range from the strength of evidence for host-lineage restriction P(*k* ≥ 0.7) in a Poisson regression. The model estimated an intercept of 1.88 (95% CI = 1.56–2.17), corresponding to approximately 6.6 species (95% CI = 4.8–8.8) when there was no evidence for host-lineage restriction (P(*k* ≥ 0.7) = 0). The estimated slope was −1.61 (95% CI = −2.32 to −0.90), meaning that a plasmid pT with strong evidence of host-lineage restriction (P(*k* ≥ 0.7) = 1) would be found in approximately 1.3 species (95% CI = 0.5–3.6). We found no effect of predicted plasmid mobility class or replicon family on the number of species.

### Conjugative backbones were mostly host-lineage restricted

Conjugation is a proposed solution to the “plasmid paradox” because it decouples plasmid survival from host survival [27]. However, ecological, physiological, and evolutionary constraints can limit a plasmid’s actual host range. Consequently, conjugative plasmids may not be widespread in a population but restricted to specific host lineages.

We quantified whether host-lineage associations varied among predicted mobility classes (*n* = 18 conjugative, *n* = 9 mobilisable, and *n* = 3 non-mobilisable; Figure 2b). Using the posterior draws of *k*, we calculated the fraction of pTs within each mobility class with *k* ≥ 0.7 (strong phylogenetic signal) or *k* ≤ 0.3 (weak phylogenetic signal). To account for uneven class sizes, we estimated these fractions via bootstrap resampling of pTs within each mobility class (*n*_boot_ = 1, 000). Then for each draw, we compared the boot-strapped fractions between mobility classes, recording the proportion of draws in which one class exhibited a stronger or weaker phylogenetic signal than another. The posterior probabilities are reported in Table 2.

**Table 2:** Posterior probabilities comparing phylogenetic signal (*k*) between plasmid mobility types. Probabilities show the fraction of posterior draws where one mobility class has higher (*k ≥* 0.7) or lower (*k ≤* 0.3) phylogenetic signal than another.

| Comparison | $k \geq 0.7$ | $k \leq 0.3$ |
| --- | --- | --- |
| Conjugative > Mobilisable | 1.000 | 0.018 |
| Conjugative > Non-mobilisable | 0.950 | 0.007 |
| Mobilisable > Non-mobilisable | 0.118 | 0.148 |

We found that conjugative pTs exhibited stronger and more consistent phylogenetic signal than both mobilisable and non-mobilisable pTs. Mobilisable pTs had values of *k* distributed approximately symmetrically around 0.5 (Supplementary Figure 1). For example, the nine pTs with the highest median *k* were F-like and B/O/K/Z-like plasmids, all predicted to be conjugative (Figure 1c–e). These pTs nonetheless exhibited clear host-lineage preferences.

Conjugative plasmids are generally longer than mobilisable or non-mobilisable plasmids. We generated a posterior sample of the association between median pT plasmid length (bp; log_10_-scaled) and *k*. We found that longer plasmids were consistently more host-lineage restricted, with a posterior median slope of 0.167 (95% CI = 0.119–0.212). In other words, when going from a small plasmid (10 kbp) to a large plasmid (100 kbp), *k* increased by approximately 0.167. Moreover, increasing plasmid length tended to strengthen host-lineage restriction more strongly for conjugative plasmids versus mobilisable plasmids (posterior median difference = 0.42, 95% CI = 0.09–0.78).

### Host-lineage restricted and conjugative backbones showed lower gain–loss ratios in ancestral state reconstructions

Ancestral state reconstruction can estimate the number of plasmid backbone independent gains and losses over evolutionary time. The ratio of gains to total events (gains + losses), or “gain-bias”, describes the propensity of a backbone to be independently and successfully acquired across lineages. A backbone with a low gain-bias is consistent with limited spread into new lineages. We reconstructed pT presence across the host phylogeny and examined gain-bias using two separate binomial regression models: one predicting gain-bias by host-lineage restriction (P(*k* ≥ 0.7)) and the other by mobility class, which allowed each factor to be interpreted marginally (see Material and Methods).

In the first model, stronger evidence of host-lineage restriction was associated with fewer independent gains across the host phylogeny (−1.89 log-odds; 95% CI = −2.22 to −1.55). In the second model, compared to conjugative pTs, mobilisable pTs showed higher gain-bias (0.27 log-odds; 95% CI = 0.07–0.46), consistent with higher realised mobility, whereas non-mobilisable plasmids showed substantially higher but more variable gain-bias (1.47 log-odds; 95% CI = 0.93–2.05), likely reflecting the small sample size (*n* = 3).

### Backbones with shared host lineages co-occurred more than expected

A pair of plasmids might co-occur due to positive interactions, because they share host lineages, because they are abundant in the population, or some combination of the three. We next sought to quantify backbone co-occurrence in a manner that distinguished contributions from shared host range and from additional interactions, while remaining robust to uneven sampling across bacterial hosts.

For each pT pair, we quantified co-occurrence along two complementary axes (Table 3). The phylogenetic contribution (Φ; -1–1) measured the extent to which co-occurrence was predicted by the relatedness of their hosts: positive values indicated that the pair was found in the same or closely related hosts, whereas negative values indicated weak or inconsistent phylogenetic structuring. The residual contribution (*R*; -1–1) captured the co-occurrence remaining after accounting for pT prevalence and host phylogeny, reflecting additional interactions not explained by host lineage. Residual values were interpreted relative to a null model that preserved each pT’s overall prevalence and lineage distribution; pairs occurring more (or less) often than expected under this model exhibited positive (or negative) residual co-occurrence. By comparing observed co-occurrence to the null expectation in this way, *R* is robust to uneven sampling across hosts and highlights genuine deviations from lineage-driven patterns (see Figure 3a–b).

**Table 3:** Interpretation of pT co-occurrence contributions. Φ quantified phylogenetic structuring, while *R* captured residual co-occurrence beyond abundance and host phylogeny.

| $\Phi$ | $R$ | Interpretation |
| --- | --- | --- |
| + | + | Co-occurrence largely explained by host phylogeny, with additional strong interactions. |
| + | – | Co-occurrence driven primarily by host phylogeny; minimal extra interactions. |
| – | + | Weak phylogenetic signal but strong additional interactions. |
| – | – | Little phylogenetic or interaction-driven co-occurrence; largely independent. |

**Figure 3:**
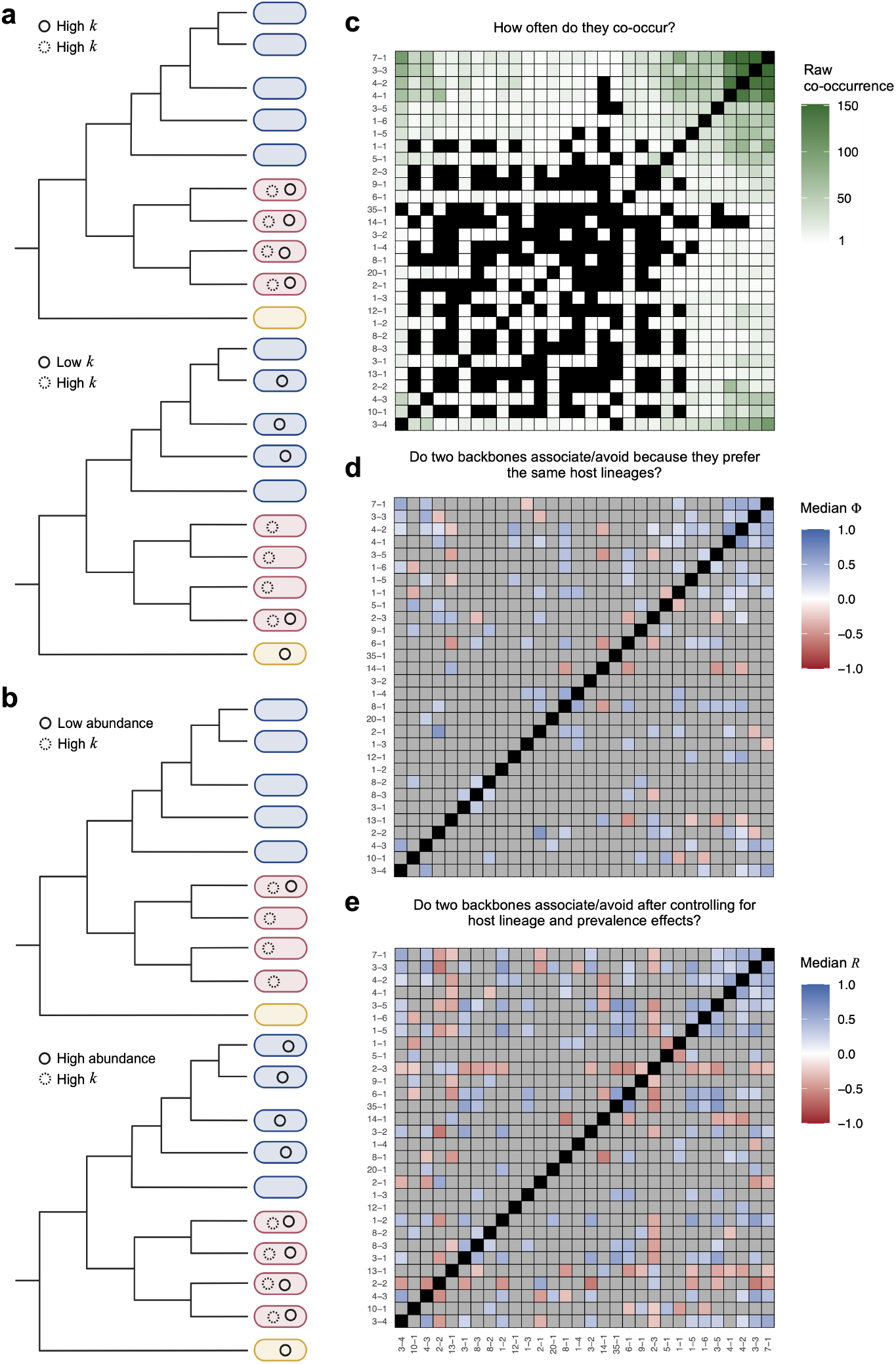
Modelling plasmid backbone co-occurrence. **(a)** Conceptual diagram illustrating how host-lineage restriction can drive plasmid co-occurrence. **(b)** Conceptual diagram illustrating how plasmid abundance can drive co-occurrence. **(c)** Raw co-occurrence counts of pT pairs across genomes, with black indicating pairs of the same pTs or pairs with zero co-occurrences. **(d)** Median Φ correlations between pTs, with red indicating positive associations, white neutral, and blue negative. Only strong correlations are shown, with the remainder in grey (see Materials and Methods). Black indicates pairs of the same pTs. **(e)** Median *R* correlations, coloured the same as Φ. Axes across panels are ordered by hierarchical clustering of raw co-occurrence counts. 19

We summarised the relationship between the two axes, finding corr(Φ, *R*) = 0.79 (95% CI = 0.66–0.89). This strong positive correlation indicated that pT pairs with similar host ranges tended to co-occur more than expected based on host phylogeny alone, whereas pT pairs with dissimilar host ranges tended to co-occur less than expected.

### Strong backbone associations or avoidances were uncommon

To assess how frequently plasmid backbones formed strong positive or negative associations, we analysed all *n* = 435 distinct pT pairs, which had a median raw co-occurrence count of 4 (range = 0–151; Figure 3c). Pairs were classified according to their phylogenetic (Φ) and residual (*R*) estimates using conservative probabilistic thresholds to identify well-supported positive or negative associations (Figure 3d–e and Table 4). In addition, to be considered positive, a pair had to be observed at least once to avoid extrapolation (see Materials and Methods).

**Table 4:** Classification of unique plasmid pT pairs (*n* = 435) based on phylogenetic (Φ) and residual (*R*) co-occurrence. For each pair, Φ represented the component of co-occurrence explained by host phylogenetic relatedness, whereas *R* represented residual co-occurrence after accounting for host phylogeny. Pairs were categorised as positive if the posterior median was positive, signed tail probability (STP) *≥* 0.95, and there was at least one co-occurrence; negative if the posterior median was negative and STP ≤ 0.95; and neutral otherwise.

| $\Phi$ class | $R$ class | Count |
| --- | --- | --- |
| Neutral | Neutral | 292 |
| Neutral | Positive | 43 |
| Neutral | Negative | 33 |
| Positive | Positive | 32 |
| Positive | Neutral | 18 |
| Negative | Negative | 16 |
| Negative | Neutral | 1 |

We found that over two-thirds (67%, 292/435) of interactions were neutral in both axes, indicating that co-occurrence was neither strongly predicted by host phylogeny nor by additional interactions. When we relaxed our thresholds for association or avoidance (from 0.95 to 0.90; see Materials and Methods), 59% (256/435) of pairs were still neutral in both axes.

Accordingly, being classified as positive in both axes was rare (7%, 32/435). For example, 10-1 was a putatively conjugative FIB-like pT that consistently carried the aerobactin and *sit* virulence operons. It showed strong host-lineage restriction: 92% (133/145) of observations were in ST95, with a median *k* = 0.83 (95% CI = 0.49–0.99) and median *σ*_*k*_ = 7.94 (95% CI = 5.66–11.44). In another example, 5-1 was a putatively conjugative B/O/K/Z-like pT carrying aminoglycoside resistance in an integron, observed 42% (80/195) of the time in ST95 (median *k* = 0.84 [95% CI = 0.45–0.99] and median *σ*_*k*_ = 1.29 [95% CI = 0.85–1.87]). Between the pair, the model estimated median Φ = 0.48 and *R* = 0.40, indicating a positive association at both host and plasmid levels, suggesting to a potentially problematic convergence of virulence and AMR determinants in ST95, a strain with a high disease burden [28].

### Host-lineage restriction of conjugative backbones drives clustering of mobilisable partners

Mobilisable plasmids exploit the conjugative machinery of co-occurring conjugative plasmids for horizontal transfer, suggesting that the distributions of conjugative and mobilisable plasmids should be related. Our model identified a clique of six mobilisable Col-like pTs (3-1, 3-3, 3-4, 4-1, 4-2, 4-3) that consistently exhibited positive co-occurrence (Φ and *R*) amongst themselves, and outwards to four conjugative F- and X-like pTs (1-5, 1-6, 8-1, 8-3; Figure 4). However, the conjugative pT themselves showed mostly neutral or negative Φ and *R* correlations.

**Figure 4:**
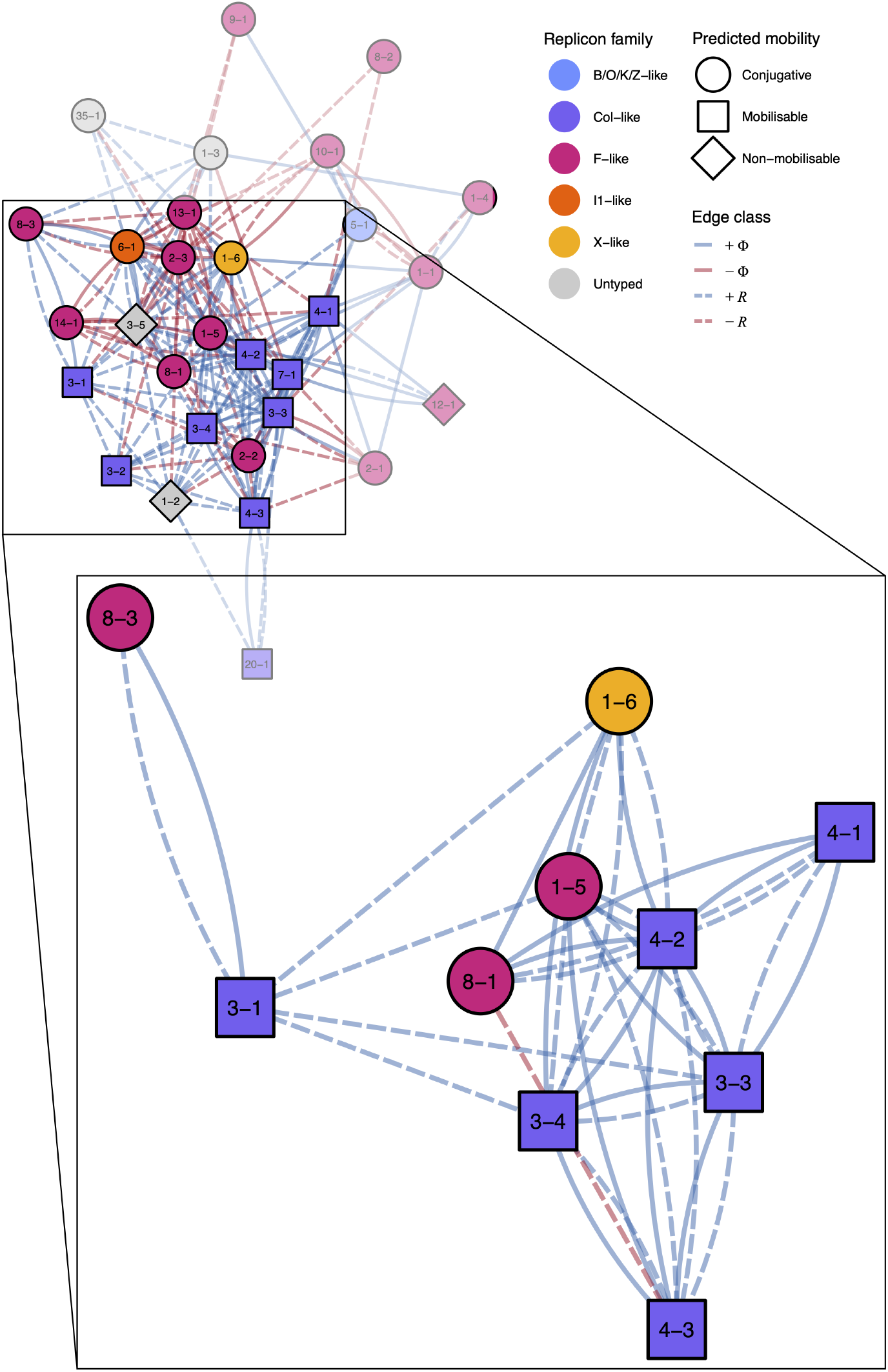
Co-occurrence network and subnetwork highlighting Φ and *R* correlations between plasmid backbones. Nodes represent pTs with shape indicating putative mobility class and fill colour indicating replicon family. Edges show positive and negative Φ and *R* correlations with neutral correlations omitted.

We hypothesised that the clustering of the mobilisable pT was driven by the host-lineage restriction of their conjugative partners. To test this, we built a mathematical model where conjugative plasmids could only transfer to a subset of host lineages, and mobilisable plasmids could only transfer via a subset of conjugative partners. This specificity reflects that not all conjugative systems mobilise all mobilisable plasmids equally [29].

We simulated host populations varying the strength of host-lineage restriction (*α* ∈ [0, 1]) and the breadth of lineages accessible to each conjugative plasmid. For each scenario, we calculated a clustering score among mobilisable plasmids as the difference between within-cluster and between-cluster co-occurrence. Our simulations recapitulated the empirical patterns (Figure 5). Mobilisable plasmids co-occurred most strongly when conjugative plasmids were restricted to a narrow set of host lineages. As conjugative plasmids could occupy more lineages, the *α* value yielding maximum clustering decreased: co-occurrence peaked at lower levels of host-lineage restriction and declined at high restriction. This occurred because broader host ranges allow compatible conjugative plasmids to overlap in more hosts even at moderate *α*, whereas strong restriction concentrates conjugative plasmids into fewer hosts than mobilisable plasmids can exploit efficiently. Clustering was minimal when host-lineage restriction was absent (*α* = 0), indicating that the observed co-occurrence arose from the combination of host-lineage restriction and overlapping conjugative partners, rather than from shared hosts alone. These patterns are consistent with host-lineage restriction being a primary driver of mobilisable plasmid clustering.

**Figure 5:**
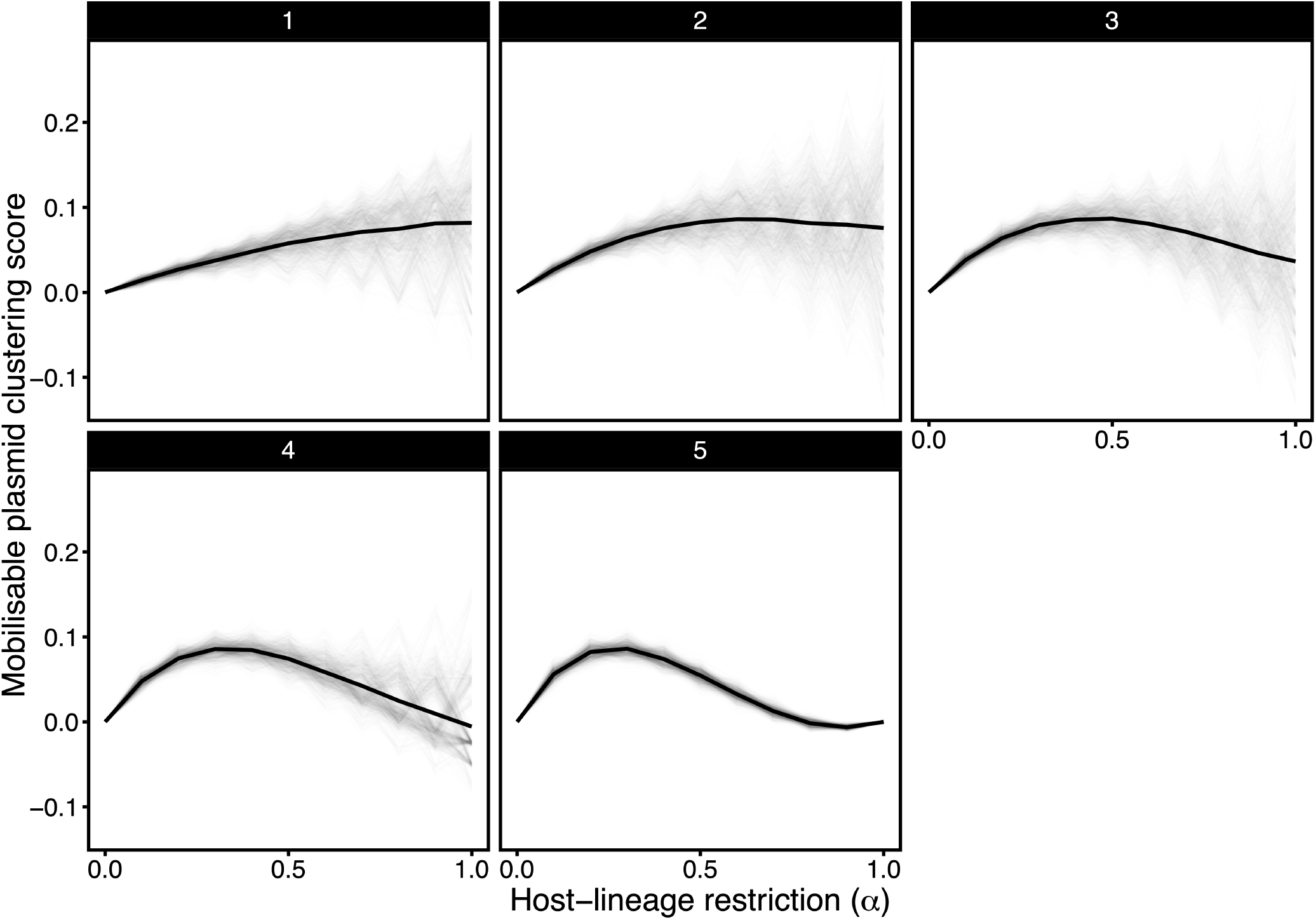
Modelling mobilisable plasmid backbone clustering. Mobilisable plasmid clustering as a function of host-lineage restriction and lineage breadth. Simulations used a mobilisation matrix in which each mobilisable plasmid could be transferred by one or more conjugative plasmids, with some overlap between mobilisable plasmids. Each panel shows the clustering signal for a given number of host lineages accessible to conjugative plasmids. Clustering scores for individual simulation replicates are shown as faint lines, and the mean clustering score across 1,000 replicates is shown in black.

## Discussion

Plasmids employ a wide repertoire of molecular mechanisms to transfer between bacterial hosts, a process central to their evolutionary persistence [27]. Here, we developed a novel Bayesian modelling framework to quantify plasmid host-lineage associations and co-occurrence while controlling for host ancestry and abundance. We show that plasmid backbones associated with *E. coli* span a continuum from host-lineage restricted to widely disseminated (Figures 1–2), uncovering two limits to realised plasmid host range: plasmid size and conjugative ability.

This raises a fundamental question: if a plasmid is largely confined to a single host lineage, what selective advantage justifies the maintenance of costly conjugative machinery? We propose that conjugation functions primarily as a mechanism for intra-lineage persistence, as opposed to inter-lineage expansion. Indeed, a key insight from previous analyses of this dataset is that the emergence of successful pathogen lineages has been driven by the acquisition of conjugative plasmids carrying genes associated with virulence and antibiotic resistance [9]. The selective advantages of these traits, coupled with the ability of plasmids to persist within a lineage, provides a parsimonious explanation for why these elements are stably maintained.

It remains unclear whether conjugative plasmids arrived in these populations already large, or accumulated length as they persisted within particular lineages. Conjugative plasmids are necessarily longer than mobilisable or non-mobilisable plasmids because they carry transfer machinery. Recent work shows that the density of restriction–modification (R-M) targets increases with plasmid size, and that smaller plasmids with fewer targets tend to have broader host ranges, presumably because they are less susceptible to R-M systems [30]. Conversely, larger plasmids, by virtue of carrying more restriction targets, may face stronger barriers to establishment in novel hosts. This suggests a mechanistic basis by which size itself could contribute to host-lineage confinement, alongside other previously discussed processes of host-specific adaptation [11, 12, 13, 14, 15]. Nonetheless, evidence for R-M systems as a size-dependent barrier stems from broad population-scale comparisons; these systems appear to be significantly less effective at the species level [31].

Conjugative systems also impose fitness costs on their hosts [32] and confer susceptibility to bacteriophages that exploit conjugative pili [33]. Together, these factors suggest that conjugative plasmids may frequently attempt transfer but fail to persist outside a restricted subset of compatible host lineages. More broadly, the limited realised mobility of conjugative plasmids may help explain the high rates of evolutionary transition from conjugative to non-conjugative plasmids reported previously, despite the apparent long-term benefits of conjugation [34].

The consequences of this restriction are visible in the structure of the within-cell plasmid communities. While a majority of backbone pairs exhibited no strong association or avoidance (Figure 3), we found that shared host range correlated strongly with excess co-occurrence. This suggests that most within-cell plasmid communities are not shaped by specific functional synergies, but rather occur opportunistically where host compatibility facilitates cohabitation. A notable exception, however, was a subnetwork of strongly associated mobilisable plasmids (Figure 4). Our simulations demonstrated that this clustering can emerge from mobilisable plasmids exploiting multiple, host-lineage restricted conjugative transfer partners (Figure 5). This is consistent with evidence that specificity between *oriT* sites and cognate relaxases constrains which partnerships are realised in practice [2].

Taken together, our results support a model in which plasmid populations are structured by nested dependencies: conjugative plasmids become confined to subsets of host lineages due to limits on establishment, while mobilisable plasmids achieve broad dissemination by exploiting multiple conjugative plasmids that occupy non-overlapping host ranges. This nested structure reconciles the apparent contradiction between low realised mobility of conjugative plasmids and the widespread distribution of mobilisable plasmids.

It is possible that the sequencing approach failed to recover some small plasmids [35]. Nonetheless, of the 30 pTs, 9 and 5 had mean lengths less than 10 kbp and 5 kbp, respectively, suggesting reasonable representation. Another limitation of the dataset is its restriction to human bloodstream infection *E. coli* isolates. However, contextualising our findings with global diversity showed that the most restricted backbones remained largely *E. coli* -specific globally, whereas less restricted backbones appeared across multiple species, indicating that observed host-lineage confinement likely reflects biological patterns rather than sampling bias.

Our findings have direct implications for the management of plasmid-mediated traits such as antimicrobial resistance and virulence. If conjugation reinforces persistence within specific host lineages, the spread of resistance genes may be more predictable and containable than previously assumed. Strategies aimed at disrupting key lineage–plasmid associations, rather than targeting plasmid transfer indiscriminately, may be more effective in limiting the emergence and persistence of high-risk plasmid genotypes. Testing this hypothesis directly represents an important direction for future work.

## Data availability

The 4,569 plasmid sequences from the original study are stably archived at https://doi.org/10.6084/m9.figshare.24302884.v1. Plasmid sample metadata and model phylogenetic signal summaries are given in Supplementary Table 1. Backbone correlation model summaries are given in Supplementary Table 2. The chromosomal phylogeny and other necessary data will be made available in the GitHub repository https://github.com/wtmatlock/plasmid-conjugation-model, which will be stably archived on Zenodo.

## Code availability

The scripts required to reproduce our analyses will be found in the GitHub repository https://github.com/wtmatlock/plasmid-conjugation-model, which will be stably archived on Zenodo.

## Acknowledgements

We especially thank Jukka Corander for help with reusing the NORM dataset and for feedback on the manuscript draft. We also thank Daniel Cazares, Edward Feil, Zamin Iqbal, Liam Shaw, and David Sünderhauf for feedback on manuscript drafts and presentations, as well as the organisers and attendees of ESCMID IMMEM XIV for the discussions that inspired this study.

---

**Supplementary Figure 1:**
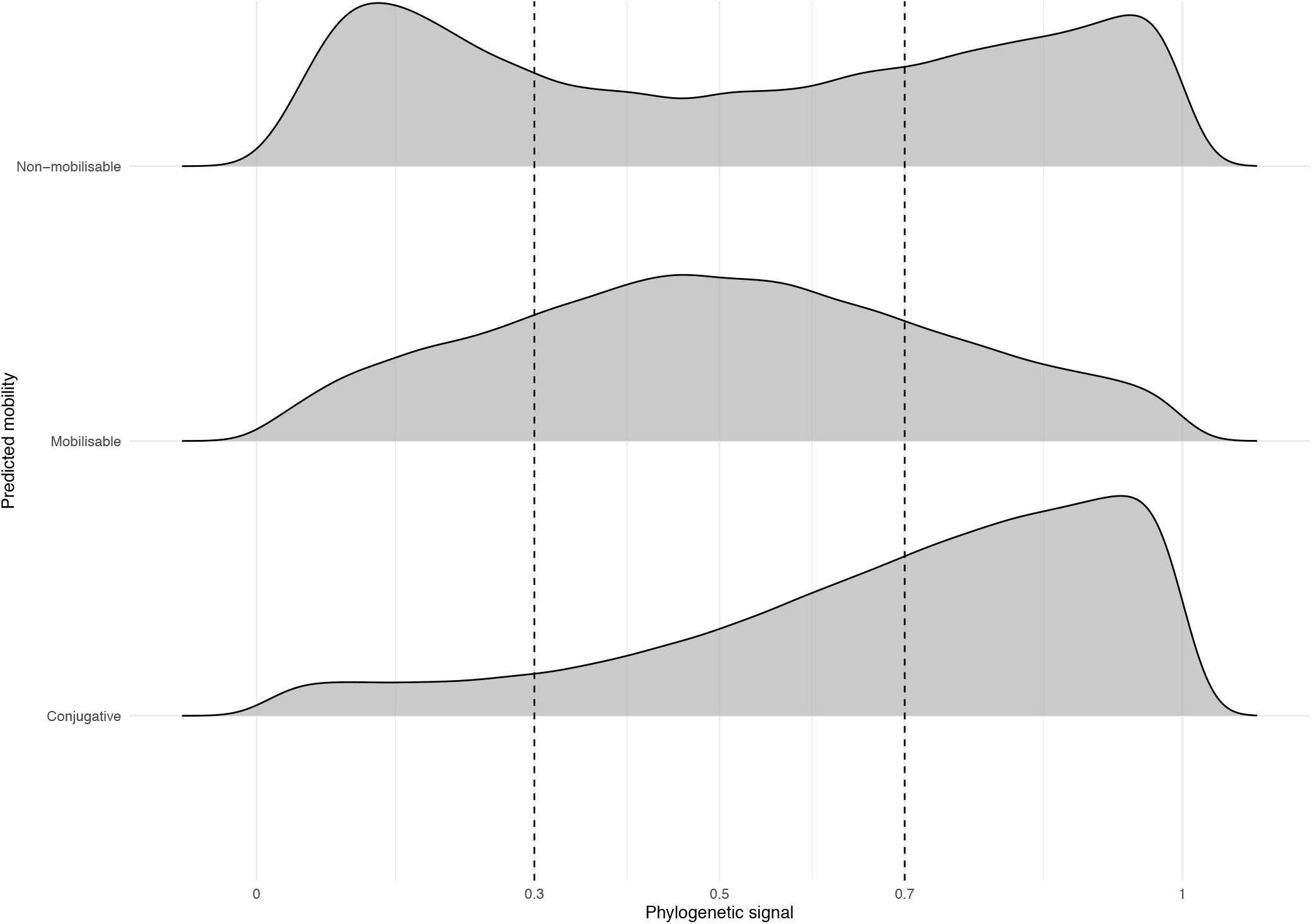
Posterior distributions of phylogenetic signal (*k*) grouped by pT predicted mobility. Of the pTs, *n* = 18 were putatively conjugative, *n* = 9 mobilisable, and *n* = 3 non-mobilisable. Values near 1 indicated strong lineage-specific structuring whereas values near 0 indicated weak or absent phylogenetic signal. Dashed lines mark thresholds for weak (*k* ≤ 0.3) and strong (*k* ≥ 0.7) phylogenetic signal.

