## Supplementary File 1 for "Conjugation structures plasmid populations through host-lineage restriction"

### Comparison to SimPhyNI

We ran SimPhyNI (v. 1.0.0) with default parameters on our dataset, modelling the presence of a pT as a binary trait.

SimPhyNI aims to detect pairs of traits that show significant association or avoidance while accounting for phylogenetic structure. It utilises ancestral state reconstruction to distinguish between genuine interactions and co-occurrence resulting from shared ancestry. Consequently, statistical significance is driven by convergent evolution: the repeated, independent gain or loss of traits across disparate branches of the phylogeny.

In total, we detected only 16 significant pT pairs (Benjamini-Yekutieli (BY)-corrected $p < 0.05$), of which 15/16 were avoidances. In all negative instances, our model showed either neutral $\Phi$ and $R$ (10/15), neutral $\Phi$ and negative $R$ (3/15), or negative $\Phi$ and negative $R$ (1/15). In the positive instance, our model showed positive $\Phi$ and positive $R$.

This outcome was expected because the power to detect positive associations in a phylogenetically-corrected model is fundamentally limited by the stability of the association. For plasmids, positive interactions often result from a single horizontal acquisition followed by stable vertical maintenance within a lineage. To SimPhyNI, this appears as a single ancestral event, providing insufficient independent data points to achieve significance. In contrast, negative interactions generate a recurrent signal of mutual exclusivity across multiple clades, providing the evidence necessary to overcome the null model.

For example, consider a 1000-tip tree where two pTs each have a 20% prevalence. Under a Poisson-based null model of independence, we expect an overlap of approximately $\mu = 1{,}000 \times 0.2 \times 0.2 = 40$ tips by chance. If these pTs were co-acquired in two disparate branches (each containing 30 tips), the resulting 60 overlaps might yield a nominal $p$-value of $P(X \geq 60) \approx 0.002$. With 30 pTs tested in total, that would set the BY-corrected threshold at $\frac{0.05}{30 \times 4} \approx 0.0004 < 0.002$, so our example fails. In contrast, observing 0 overlaps, $P(X = 0) \approx 0$, which passes the threshold.
